# *Wolbachia* and its pWCP plasmid show differential dynamics during the development of *Culex* mosquitoes

**DOI:** 10.1101/2025.01.03.631243

**Authors:** Alice Brunner, Camille Gauliard, Jordan Tutagata, Seth Bordenstein, Sarah Bordenstein, Blandine Trouche, Julie Reveillaud

**Affiliations:** Mivegec, Université de Montpellier, INRAE, CNRS, IRD, Montpellier, France; Department of Department of Biology, Eberly College of Science, Pennsylvania State University, University Park, PA, USA

**Keywords:** Mosquitoes, Wolbachia, Mobile genetic elements, plasmid, Life cycle

## Abstract

Mosquitoes are major vectors of pathogens such as arboviruses and parasites, causing significant health impacts each year. *Wolbachia*, an intracellular bacterium widely distributed among arthropods, represents a promising vector control solution. This bacterium can reduce the transmission of dengue, Zika and chikungunya arboviruses and manipulate the reproduction of its host through its prophage WO. Although research on the *Wolbachia* mobilome primarily focuses on WO and the phenotypes it induces, the function of *Wolbachia* plasmid pWCP, recently discovered and reported to be strikingly conserved worldwide, remains unknown. In this study, we analyzed the presence and abundance of pWCP as well as *Wolbachia* in two different species of *Culex* mosquitoes, one of the most widespread genera in the world and a vector of numerous diseases. We compared relative densities of the bacterium and its mobile genetic element in *Culex pipiens molestus* and *Culex quinquefasciatus*, a facultatively autogenous and a anautogenous species, respectively, throughout their development from larval stage L1 to adult individual specimen using quantitative PCR. Our results suggest that 2-5 copies of pWCP occur in *Wolbachia* cells on average, and the plasmid co-replicates with *Wolbachia* cells. Moreover, *Wolbachia* and pWCP exhibit differential levels of abundance at specific development stages throughout the mosquito’s life cycle in each species. These findings indicate important, and likely beneficial, roles for the plasmid in the bacterium’s biology in different mosquito species as well as complex interaction dynamics between *Wolbachia* and its host during its life cycle.

**Importance:** Mosquitoes of the *Culex* genus are critical vectors for numerous diseases, causing significant public health concerns. The intracellular bacterium *Wolbachia* has emerged as a promising vector control solution due to its ability to interfere with pathogen transmission and manipulate mosquito reproduction. However, unlike the extensively studied WO phage, the biological significance and function of *Wolbachia’s* pWCP plasmid, a recently discovered and strikingly conserved mobile genetic element in *Culex* species, remains unknown. This study investigates the developmental dynamics of pWCP and *Wolbachia* in two *Culex* mosquito species, *Culex pipiens molestus* and *Culex quinquefasciatus* across their life cycle. In general, the abundance levels of *Wolbachia* and the plasmid were found to vary across life stages and differ between the two species. However, a relatively small number of pWCP copies were observed per *Wolbachia* cell, together with a co-replication of the plasmid with the bacterium for most developmental stages. Altogether, these findings suggest a likely beneficial and non-parasitic role for pWCP in *Wolbachia*’s biology, that may contribute to the intricate interactions between the bacterium and its mosquito hosts.

## Introduction

Mobile genetic elements (MGEs) are essential drivers of genomic plasticity and adaptation in organisms, notably by facilitating horizontal gene transfer (Marimuthu et al., 2022; Tria & Martin, 2021; Weisberg & Chang, 2023). They include diverse entities such as insertion sequences (IS), transposons (Tn), integrons (In), phages, and plasmids, which have the ability to move or spread within the same genome or between different cells, thereby altering the genetic structure of their hosts (Fitzgerald et al., 2021; Sengupta & Austin, 2011; Siguier et al., 2014; The et al., 2016). Through their various mechanisms of propagation, such as conjugation (mediated by plasmids and integrative conjugative elements), transduction (mediated by phages), and transformation (uptake of extracellular DNA), these MGEs actively contribute to the acquisition of new functions and the rapid evolution of microbial communities (Lerminiaux & Cameron, 2019; Partridge et al., 2018).

Phages and plasmids, as major categories of MGEs, have a particularly significant impact on bacterial evolution (Mavrich & Hatfull, 2017; Wein & Dagan, 2020). In addition to their core genes, these MGEs typically carry accessory genes that provide selective advantages to their host cells, such as antibiotic resistance, virulence factors, or unusual metabolic pathways (Dedrick et al., 2021; Frost et al., 2005; Richardson et al., 2018). Plasmids, which are present in all domains of life (Kazlauskas et al., 2019), generally exist as circular extrachromosomal DNA and replicate independently of the bacterial chromosomal DNA (del Solar et al., 1998). To coexist stably with their hosts and minimize metabolic burden, plasmids must regulate their replication so that their copy number remains consistent within a given host and under defined growth conditions (Del Solar & Espinosa, 2000). However, recent studies show that modulating plasmid copy numbers can directly influence bacterial growth and have significant impacts on the metabolic costs borne by the host (Rouches et al., 2022). For example, the variability in plasmid copy number can result in varied effects on the stability of replication, plasmid loss, or even impose a metabolic burden, thereby influencing bacterial interactions within the host cell (Rouches et al., 2022).

In addition, recent advances in metagenomics, in particular state-of-the-art assembly and binning approaches are shedding light on the diversity and dynamics of plasmids that lack identifiable traits or detectable host markers (Attéré et al., 2017; Challacombe et al., 2017; Yu et al., 2024). For example, the cryptic plasmid pBI143 of *Bacteroides fragilis*, recently discovered through metagenomic studies of the human gut microbiota, was found as one of the most abundant elements in this ecosystem (Fogarty et al., 2024). Although its precise function remains unknown, the relative copy number of the element increases during stress such as Inflammatory Bowel Disease (Fogarty et al., 2024). Similarly, the first identified plasmid of *Wolbachia*, named pWCP for *Wolbachia* plasmid in *Culex pipiens*, was found as highly conserved in *Culex* spp., and present worldwide (Ghousein et al., 2023). Data showed a rather stable copy number of 4-5 in the ovaries of adult females worldwide (Ghousein et al., 2023), yet with some inter-individual variations possibly reflecting distinct physiological states of the mosquito hots (Ghousein et al., 2023). Overall, the discovery of pWCP could open new possibilities for effective genome-editing strategies in a bacterium that has, until now, been resistant to genetic modification.

*Wolbachia* was first identified in 1924 (Hertig & Wolbach, 1924) and belongs to the order *Rickettsiales* (Alphaproteobacteria). The main evolutionary lineages of *Wolbachia* are referred to as “supergroups” and are designated by letters (A-F, H-K and S) (Zhou et al., 1998). Supergroups A and B are found in many terrestrial arthropods (Gerth et al., 2014) and infect over 70% of these arthropods (Jeyaprakash & Hoy, 2000), including 50% of insect species (Fallon, 2021). Its prevalence and frequency within insect populations are explained by its ability to manipulate host reproduction (Werren et al., 2008) and possibly by nutritional mutualism (Newton & Rice, 2020).

*Wolbachia* can induce various reproductive manipulation phenotypes, notably cytoplasmic incompatibility (CI) (Kaur et al., 2021; Shropshire et al., 2020). CI causes sterility in crosses between infected males and uninfected females or between individuals infected with different and incompatible *Wolbachia* strains, facilitating the establishment of infected populations (Ant et al., 2020; Dutton & Sinkins, 2004). This phenomenon is used in biocontrol programs, such as the World Mosquito Program, based on the release of *Aedes aegypti* mosquitoes infected with *Wolbachia* strains to fight diseases like dengue and Zika (Caputo et al., 2020; Chambers et al., 2011; O’Neill, 2018). The effectiveness of this method is enhanced by *Wolbachia*’s ability to block pathogen transmission in mosquitoes (Ahmed et al., 2016; Ant et al., 2020; Liang et al., 2020; Moran et al., 2008). Mosquitoes trans-infected with *Wolbachia* show varying levels of resistance to dengue, Zika, and chikungunya viruses, with viral inhibition often linked to *Wolbachia* density in somatic tissues (Liang et al., 2020).

CI is mediated by *cifA* and *cifB* genes located in the WO prophage of Wolbachia (LePage et al., 2017). Recent findings show that plasmids in *Wolbachia* and related bacteria, such as *Rickettsia*, carry *cif* gene homologs, indicating a potential role of horizontal plasmid transfer in CI acquisition (Martinez et al., 2022; Owashi et al., 2024). Others plasmids, notably pWALBA1 and pWALBA2 in *Wolbachia* of *Aedes albopictus* (*w*AlbA) showed features rather similar to pWCP (Martinez et al., 2022) with many unknown genes. The mobile genetic element pWCP comprises 9.23 kbp encoding 14 genes (Reveillaud and Bordenstein et al., 2019). Of these, seven have known functions, including some involved in DNA replication (*DnaB*-like helicase), plasmid partitioning (*ParA*-like), toxin-antitoxin stability systems (*RelBE* loci), and a transposable element (IS110-family), which collectively suggest a role in plasmid maintenance and mobility. The remaining seven genes are of unknown function (Reveillaud and Bordenstein et al., 2019). It is possible that one specific or a set of pWCP genes play a key role for the bacterium and the mosquito holobiont as a whole, particularly during physiological changes or insect metamorphosis, which might influence pWCP copy numbers in certain conditions. This is observed in the symbiont *Buchnera aphidicola* of aphids, where the copy number of the leucine biosynthesis plasmid increases under amino acid deprivation in the host (Viñuelas et al., 2011).

In this study, we explore the relative abundance of plasmid pWCP and *Wolbachia* itself throughout the development of two mosquito species of the *Culex* genus, *Culex pipiens molestus* and *Culex quinquefasciatus,* a facultatively autogenous and anautogenous species, respectively. Mosquitoes were analyzed from the first larval stage to the fourth larval stage, pupae and adults (male and female) stages using quantitative PCR to determine the variability or stability of the plasmid and *Wolbachia* across the different stages of development.

## Materials and Methods

### Mosquito Rearing and Sampling

We maintained *the Culex pipiens molestus* (Celestine strain, autogenous, originally from Montpellier, reared in the laboratory since October 2023, 5^th^ generation) and *Culex quinquefasciatus* (SLAB strain, initially isolated from the San Joaquin Valley, California, USA, in the 1960s, ca. 96^th^ generation in our laboratory) at 27°C with 70% relative humidity and a 12h-12h light-dark cycle. We provided 10% sugar water weekly in Erlenmeyer flasks for both strains.

The SLAB strain, being anautogenous, additionally received a monthly blood meal from canaries (*Serinus canaria*) restrained in the mosquito cages. We reared larvae from both species in insectary trays (30 x 20 x 6 cm) filled with 1L osmosis water at a density of approximately 200 larvae per tray. The larval diet consisted of a mixture of 1/3 Tetramin fish food (Tetra, Germany) and 2/3 rabbit pellets (Versele-Laga, Belgium).

We collected 25 individuals at each developmental stage (L1, L2, L3, L4, pupa, adult female, adult male) from each species (n=175) from two trays per species (4 trays in total). We sampled larvae every 48 hours during stages L1-L4 to ensure distinct developmental stages. For adult females, we collected individuals 24h after emergence, ensuring they had not taken a blood meal. We preserved each sampled individual in 100 µL of sterile phosphate-buffered saline (PBS) and stored them at −20°C.

### DNA Extraction and quantitative Real-Time PCR

We extracted DNA from each individual using the Qiagen “DNeasy Blood and Tissue Kit” (Qiagen, Hilden, Germany) following the manufacturer’s instructions and quantified it using the Qubit dsDNA HS Assay Kit (Thermo Fisher Scientific, Waltham, MA, USA, Invitrogen ™). We performed quantitative PCR (qPCR) on all individuals of both species (*Cx. quinquefasciatus* and *Cx. pipiens molestus*). All qPCRs (45 cycles of 95°C for 10s, 58°C for 20s, and 65°C for 20s) were performed on the Lightcycler LC480 real-time PCR instrument (Roche diagnostics, Mannheim, Germany) and the SensiFAST SYBR No-ROX Kit (Bioline, Meridian Bioscience, London, UK), according to the manufacturer’s instructions. We carried out each qPCR reaction in duplicate using 6µL mix (2.5µL of SYBR No-ROX, 1µL of DNA, each primer at 0.6µM, adjusted to a total volume of 6µL with nuclease-free water (Qiagen, Hilden, Germany).

We used specific primers (**Supplementary Table 1**) to target the *Wolbachia* surface protein gene (*wsp*), plasmid pWCP gene (*GP11*), and the mosquito acetylcholinesterase gene (*ace2*), all present in single copy. Serial dilutions of a synthetic plasmid (Eurofins Scientific SE, Luxembourg) including the genes *GP11*, *wsp*, and *ace2* were placed in triplicate on each qPCR plate (4 serial dilutions per plate) for each gene. These served as internal controls to generate standard curves for each gene and each qPCR plate, mitigating any potential bias from variations between qPCR plates. Fluorescence data were analyzed with the LightCycler480 software, which converted Ct values into concentrations (ng/µL) based on the standard curves. Samples with Cp values greater than 30 were excluded from the analysis.

We determined the relative proportions of each gene by normalizing the calculated concentrations of *wsp* and *GP11* genes to gene *ace2*, allowing to account for variations in DNA quantities between samples.

### Statistical Analyses

We performed all statistical analyses and data visualizations using R software (R Core Team, version 4.3.1, Vienna, Austria, 2023). We analyzed data from the two mosquito species independently, with each undergoing identical analyses. Linear models were used to investigate plasmid quantity (*GP11/wsp* ratio) and *Wolbachia* quantity (*wsp/ace2* ratio) across developmental stages. Generalized linear mixed models (GLM, lme4 package) (Bates et al., 2003) were applied. Model assumptions (normality and homoscedasticity of residuals) were verified using the DHARMa (Hartig, 2016) and Performance (Lüdecke et al., 2021) packages. Parameters were analyzed under a normal distribution, and pairwise comparison tests were conducted to compare plasmid and *Wolbachia* quantities between stages.

We performed correlation analyses to examine the relationship between *Wolbachia* quantities and plasmid pWCP quantities at each developmental stage and for each individual. The choice between Pearson and Spearman correlation tests was determined by testing the raw data for normality within each developmental stage. We then applied either Student’s t-tests (for normally distributed data) or Wilcoxon rank-sum tests (for non-normal data) to compare *Wolbachia* quantities between species at each developmental stage. Of note, correlation analyses are calculated on an individual basis while comparison of dynamic analyses for *Wolbachia* and its plasmid pWCP focus on the developmental stage level as a whole.

## Data availability

Raw qPCR data for *Culex pipiens molestus* and *Culex quinquefasciatus* are available in Supplementary Table 2 and 3, respectively. Raw *Wolbachia* and pWCP quantities for species comparison are available in Supplementary Table 4. A reproducible bioinformatics workflow (R code) used for this study is available in Supplementary Note 1.

## Results

### Wolbachia and pWCP plasmid abundance in Culex pipiens molestus

We quantified the dynamics of *Wolbachia* and pWCP plasmid across the different developmental stages of *Culex pipiens molestus* (**Figure 1A-C**). For *Wolbachia* quantities (**Figure 1.A**), we observed an average bacterial copy number (*wsp/ace2* ratio) ranging from 0.165 ± 0.213 (at the L3 stage) to 0.805 ± 0.474 (in adult females), with a significant increase at the pupal and adult female stages compared to other stages (p < 0.0001). For pWCP counts, the average ratio of the copy number of the plasmid by *Wolbachia* cell ranged from 2.69 ± 1.65 (in adult females) to 5.22 ± 1.48 (at the L3 larval stage), indicating that there are multiple copies of the plasmid in each cell. Pairwise comparisons adjusted using the Bonferroni method revealed a significant increase in plasmid copy number by *Wolbachia* cell at the L3 stage, notably compared to adults females (SE = 0.539; p = 0.0002) and to the pupal stage (SE = 0.527; p = 0.0005) (**Figure 1.B**).

**Figure 1:**
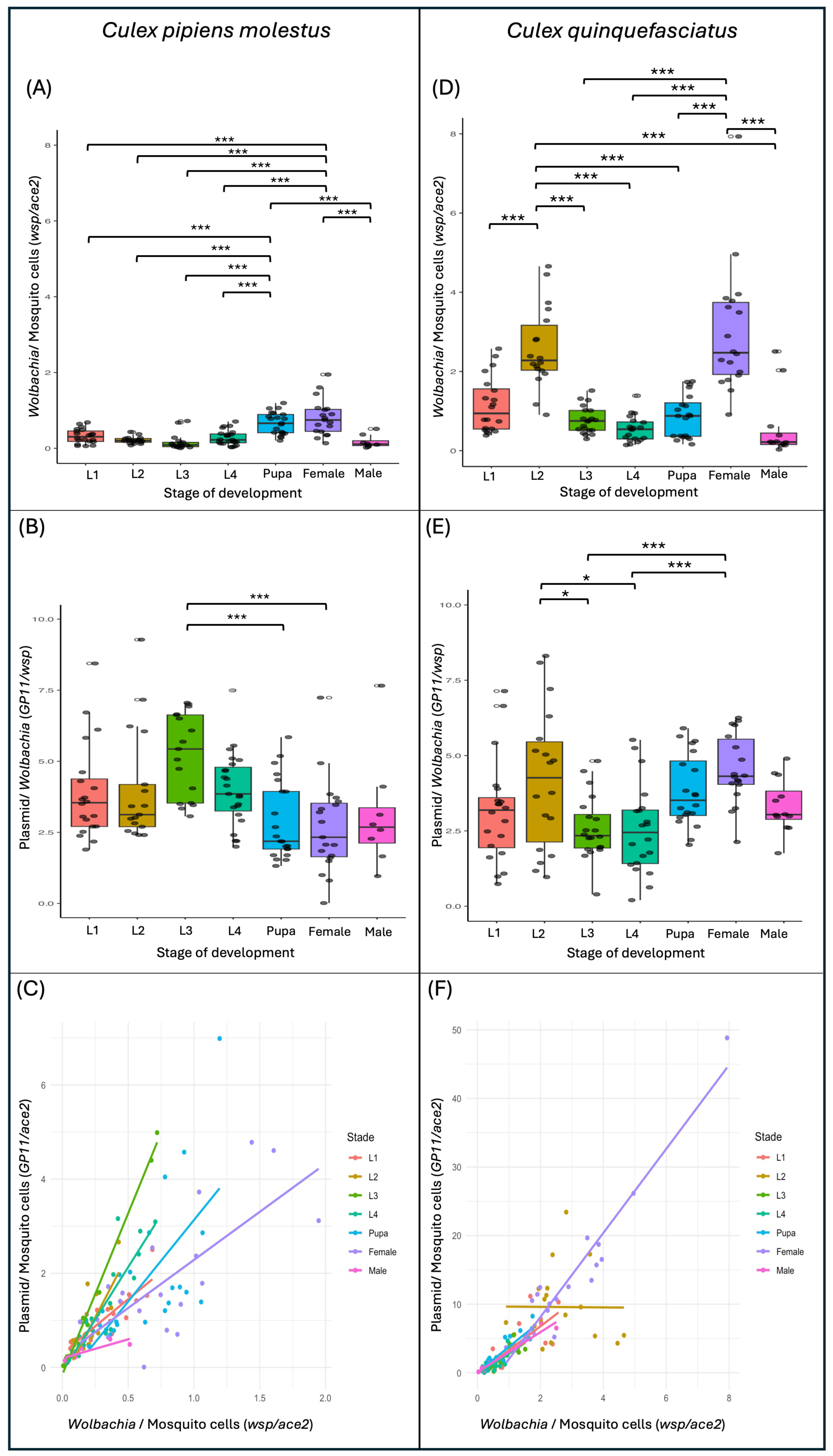
**Dynamics of plasmid and *Wolbachia* quantities during the development of *Culex pipiens molestus* and *Culex quinquefasciatus* mosquitoes.** The relative quantities of *Wolbachia* (*wsp* gene) are shown in (A) and (D) for *Culex pipiens molestus* and *Culex quinquefasciatus*, respectively, while the ones of the plasmid pWCP (*GP11*) are shown in (B) and (E) for each species. Correlation analyses between *Wolbachia* and plasmid quantities are represented in (C) for *Culex pipiens molestus* and in (F) for *Culex quinquefasciatus*. Boxplots indicate the median and interquartile range of the data. While gray dots represent individual values while white circle indicate outliers identified in the dataset. * Indicates significance < 0.05, ** indicates significance < 0.01, and *** indicates significance < 0.001. In (C) and (F), all correlations are significant (<0.001), except for the L2 stage in (F).

Notably, we observed a significant positive correlation between the densities of pWCP and *Wolbachia* (p < 0.01) (**Figure 1.C**). A particularly strong slope and correlation were observed at the L3 larval stage (R = 0.995; p = 2.28E-16, Pearson correlation), indicating a higher number of plasmid copies per *Wolbachia* at this stage compared to the other developmental stages. Overall, slope values varied according to the developmental stage, indicating that plasmid copy numbers may change throughout development (**Figure 1.C**), but overall the plasmid co-replicates with the *Wolbachia* cells.

### Wolbachia and pWCP plasmid abundance in Culex quinquefasciatus

In *Culex quinquefasciatus* (**Figure 1**), *Wolbachia* copy number ranged from 0.545 ± 0.323 (at the L4 stage) to 2.99 ± 1.62 (in females), with a significant increase at the L2 and female stages compared to the other stages (p < 0.0001) (**Figure 1.D**). For the pWCP plasmid, copy number ranged from 2.54 ± 1.48 (at the L4 stage) to 4.55 ± 1.16 (in females) (**Figure 1.E**), which are similar to that of *Culex pipiens molestus* and indicative of multiple plasmid copies per *Wolbachia* cell. Adjusted pairwise comparisons showed that plasmid quantities were higher in adults females and L2, significantly compared to the L3 (SE = 0.485; p = 0.0021; SE = 0.485; p = 0.0258) and L4 stages (SE = 0.485; p = 0.0013; SE = 0.485; p = 0.0170), respectively.

Correlation analysis between *Wolbachia* and plasmid densities revealed similarly positive correlations for all but the L2 larval development stage of *Culex quinquefasciatus* (**Figure 1.F**) (R = −0.007; p = 0.97). At this particular stage, some individuals displayed a marked increase in *Wolbachia* without a concomitant increase in plasmid copies, unlike at other stages. Consequently, these results suggest that the plasmid co-replicates with the *Wolbachia* cells except during this larval stage.

### Comparison of Wolbachia quantities between Culex pipiens molestus and Culex quinquefasciatus

Next, we investigated potential differences in *Wolbachia* quantities (*wsp*/*ace2* ratio) between *Culex pipiens molestus* and *Culex quinquefasciatus*. Results revealed significant differences between the two species at most stages. We observed a highly significant difference in *Wolbachia* density between species at the female (W = 11, p = 2.21 × 10^−8^), L1 (W = 30, p = 4.07 × 10^−7^), L2 (t = −9.54, p = 5.19 × 10^−11^), L3 (W = 18, p = 2.01 × 10^−7^), and L4 (t = −3.35, p = 0.0017) stages, with *Culex quinquefasciatus* consistently showing higher densities than *Culex pipiens molestus*. However, no significant differences in *Wolbachia* density were observed at the male (p = 0.0691) or pupal (p = 0.3862) stages.

## Discussion

This study investigates the presence and dynamics of *Wolbachia* and pWCP plasmid across the developmental stages of *Culex pipiens molestus* and *Culex quinquefasciatus*. Our results show that both *Wolbachia* and its pWCP plasmid are present at all developmental stages, and for nearly all stages, the plasmid co-replicates with *Wolbachia* cells with roughly 2-5 copies per bacteria. Minor variations in counts or densities may reflect stage-specific as well as species-specific dynamics. Furthermore, although variations in the copy number of pWCP plasmid were mostly correlating with bacterial density, exceptions at the L2 larval stage of one species suggested a desynchronization of replication and complex interactions between *Wolbachia* and its associated mobile genetic element.

We observed differences in *Wolbachia* quantities between individuals at the same developmental stage within the same species. However, these variations were relatively minor, with *Wolbachia* levels varying up to 2- to 3-fold between individuals within the same stage. In contrast, variations observed in natural insect populations such as *Drosophila* can be much more substantial, reaching up to 20,000-fold differences between certain individuals (Unckless et al., 2009). This low variability in our laboratory colonies suggests a relative stability of *Wolbachia* levels under controlled environmental conditions. Nonetheless, differences persist despite these stable conditions, reflecting the inherent variability of *Wolbachia* densities in infected hosts. Overall, intra-stage variations remained nevertheless lower than inter-stage variations.

The distinct *Wolbachia* dynamics observed between the two species—with increased bacterial abundance starting at the pupal and adult female stages in *Culex pipiens molestus*, and at the adult female stage in *Culex quinquefasciatus*—may be linked to differences in their reproductive biology. *Culex pipiens molestus* is an autogenous species capable of reproducing without a prior blood meal, unlike *Culex quinquefasciatus*, which is anautogenous and requires a blood meal to initiate ovarian development. In an autogenous species like *Culex pipiens molestus*, the previtellogenic phase, which allows follicles to develop and become competent for fertilization, may begin earlier, potentially at the pupal stage (Clements, 1999). The release of juvenile hormone, necessary to progress beyond stage 1 of previtellogenic development and which induces an increase in bacterial abundance in the ovaries (Fallon, 2021), may occur before emergence in autogenous species, unlike in anautogenous species, where it is released only after the first blood meal. Since *Wolbachia* is highly concentrated in the ovaries (Hague et al., 2024), the early development of ovaries in this species could explain the bacterium’s increase observed at the pupal stage. A previous study reported an increase in *Wolbachia* levels at the adult female stage compared to the larval stages in the SLAB strain (*Culex quinquefasciatus*), although only the L4 stage was sampled (Berticat et al., 2002). A similar increase between the L4 and female stages was observed in the SLAB strain in this study, but an additional increase in *Wolbachia* was also detected at the L2 larval stage. These results suggest either a stress-induced response or a physiological change occurring at this developmental stage that would induce a sudden increase in the intracellular bacterium, thought the exact causes and triggers remain to be determined. In both species, *Wolbachia* quantities were nevertheless higher in females than in males, a result consistent with previous observations in other species such as *Drosophila simulans* (Bourtzis et al., 1998; Rousset et al., 1999) and *Aedes albopictus* (Dobson et al., 1999). Although *Wolbachia* are present in various tissues, it is primarily concentrated in the ovaries. The higher infection load in females can therefore be attributed to the significantly larger size of ovaries compared to testes.

When examining *Wolbachia* and plasmid quantities for each individual, we observed variations in pWCP plasmid copy number along the mosquito life cycles, as well as differences between the two species. In *Culex pipiens molestus*, variations in the pWCP plasmid were positively correlated with variations in *Wolbachia* levels at all developmental stages, indicating that when *Wolbachia* increase, the plasmid copy number also increases (**Figure 1.C**). This observed correlation differs from the previously reported inverse correlation between temperate phage WO and *Wolbachia* in *Nasonia* parasitoid wasps (Bordenstein et al., 2006) whereby the phage can lyse *Wolbachia* cells during is replication stages to potentially invade new *Wolbachia* cells. A stronger correlation was observed at the L3 larval stage, suggesting a higher number of plasmid copies per *Wolbachia* at this specific stage. This increase in plasmid copy number may suggest a L3 stage-specific function of the plasmid in this species. Nevertheless, the plasmid overall co-replicates with the *Wolbachia* cells, suggesting a non-parasitic mobile element in this species.

In *Culex quinquefasciatus*, a positive correlation between *Wolbachia* and pWCP copy numbers was observed across all developmental stages, except at the L2 stage, where a marked lack of correlation between *Wolbachia* and pWCP densities was detected (**Figure 1.F**). These results suggest a specific stress or physiological change at L2 stage, and that a notable increase in the intracellular bacterium load might cause at least a temporary desynchronization of plasmid replication. It would be pertinent to sample this L2 stage at different time points in *Culex quinquefasciatus* with higher sample sizes to see if resynchronization of plasmid and *Wolbachia* replication occurs; this could better pinpoint the potential source of stress. It is possible that some *Wolbachia* cells may replicate too quickly for the plasmid to replicate and/or segregate properly, leading to the loss or to a lower number of the mobile genetic element in part of the *Wolbachia* population. However, the presence of two *RelBE* toxin–antitoxin (TA) systems in the pWCP plasmid may promote the stability of the mobile element by killing cells that lose the antitoxin component after segregation (Yarmolinsky, 1995).

Generally, we observed that *Wolbachia* and pWCP plasmid quantities were consistently higher in *Culex quinquefasciatus* than in the autogenous *Culex pipiens molestus*. This observation aligns with previous studies that have shown interspecific differences between autogenous and anautogenous species in terms of *Wolbachia* densities, particularly in the testes, where strains such as Maclo (*Culex quinquefasciatus*) exhibit higher *Wolbachia* densities than strains such as Tunis (*Culex pipiens molestus*) (Duron et al., 2007). However, the density and variability differences observed between the two *Culex* species may result not only from their different life strategies (autogenous vs. anautogenous) but also from their distinct geographic origins (North America vs. France herein), as well as genetic differences between the *Wolbachia* groups in *Culex pipiens* (*w*Pip) to which they belong. The SLAB strain (*w*Pip group III) (Dumas et al., 2013) and the Celestine strain (*w*Pip group I or III according to Dumas et al., 2013) show allelic differences in several genes (Atyame et al., 2011), which could explain the variations in *Wolbachia* population dynamics and the abundance differences observed between the two species.

## Conclusion

The study of several mosquito species with distinct life cycles highlighted complex interactions and behaviors between pWCP and its hosts. Overall, variation of plasmid copy number per *Wolbachia* cell suggest largely stable multi-copy replication of the plasmid during mosquito development. The plasmid’s copy number, stability, and transfer frequency could be influenced by plasmid gene and regulatory elements, which appear to be highly conserved across species, but also by host factors. The latter could affect the persistence and evolutionary dynamics of the plasmid within a population in different ways. Further studies to attempt localizing plasmid accumulation in *Wolbachia* from different tissues (ovaries, midgut, etc.) using state of the art microscopy tools, as well as expression studies of the plasmid’s different genes, could help investigating the variations of the plasmid at finer scale and better define its potential role in mosquito species.

## Aknowledgments

We thank Angélique Porciani for helpful discussion on statistical analyses. This work was supported by the ERC RosaLind Starting Grant “948135” to JR. We thank the Vectopole platform (IRD, Montpellier, France) for providing technical support and for the rearing and maintenance of the mosquito populations. The Vectopole is a platform of the ‘Vectopole Sud’ Network and is part of the LabEx CeMEB (ANR-10-LABX-04-01).

## Authors contributions

AB designed and performed the experiments, analysed the data, prepared the figures and wrote the manuscript. CG and JT contributed to the experiment design, coordinated laboratory experiments and wrote the manuscript. Sarah Bordenstein and Seth Bordenstein participated in the conception of this study and wrote the manuscript. BT coordinated data analysis and wrote the manuscript. JR conceived and coordinated this study, and wrote the manuscript.

## References

Ahmed, M. Z., Breinholt, J. W., & Kawahara, A. Y. (2016). Evidence for common horizontal transmission of *Wolbachia* among butterflies and moths. BMC Evolutionary Biology, 16(1), 118. 10.1186/s12862-016-0660-x

Ant, T. H., Herd, C., Louis, F., Failloux, A. B., & Sinkins, S. P. (2020). *Wolbachia* transinfections in *Culex quinquefasciatus* generate cytoplasmic incompatibility. Insect Molecular Biology, 29(1), 1–8. 10.1111/imb.12604

Armoo, S., Doyle, S. R., Osei-Atweneboana, M. Y., & Grant, W. N. (2017). Significant heterogeneity in *Wolbachia* copy number within and between populations of *Onchocerca volvulus*. Parasites & Vectors, 10(1), 188. 10.1186/s13071-017-2126-4

Attéré, S. A., Vincent, A. T., Paccaud, M., Frenette, M., & Charette, S. J. (2017). The Role for the Small Cryptic Plasmids As Moldable Vectors for Genetic Innovation in Aeromonas salmonicida subsp. Salmonicida. Frontiers in Genetics, 8. 10.3389/fgene.2017.00211

Atyame, C. M., Delsuc, F., Pasteur, N., Weill, M., & Duron, O. (2011). Diversification of *Wolbachia* Endosymbiont in the *Culex pipiens* Mosquito. Molecular Biology and Evolution, 28(10), 2761–2772. 10.1093/molbev/msr083

Bates, D., Maechler, M., Bolker, B., & Walker, S. (2003). *lme4 : Linear Mixed-EZects Models using « Eigen » and S4* (p. 1.1–35.5) [Jeu de données]. 10.32614/CRAN.package.lme4

Berticat, C., Rousset, F., Raymond, M., Berthomieu, A., & Weill, M. (2002). High *Wolbachia* density in insecticide–resistant mosquitoes. Proceedings of the Royal Society of London. Series B: Biological Sciences, 269(1498), 1413–1416. 10.1098/rspb.2002.2022

Bordenstein, S. R., Marshall, M. L., Fry, A. J., Kim, U., & Wernegreen, J. J. (2006). The Tripartite Associations between Bacteriophage, *Wolbachia*, and Arthropods. PLOS Pathogens, 2(5), e43. 10.1371/journal.ppat.0020043

Bourtzis, K., Dobson, S. L., Braig, H. R., & O’Neill, S. L. (1998). Rescuing Wolbachia have been overlooked⃛. Nature, 391(6670), 852–853. 10.1038/36017

Caputo, B., Moretti, R., Manica, M., Serini, P., Lampazzi, E., Bonanni, M., Fabbri, G., Pichler, V., della Torre, A., & Calvitti, M. (2020). A bacterium against the tiger : Preliminary evidence of fertility reduction after release of Aedes albopictus males with manipulated *Wolbachia* infection in an Italian urban area. Pest Management Science, 76(4), 1324–1332. 10.1002/ps.5643

Caragata, E. P., Dutra, H. L. C., & Moreira, L. A. (s. d.). Inhibition of Zika virus by *Wolbachia* in *Aedes aegypti*. Microbial Cell, 3(7), 293–295. 10.15698/mic2016.07.513

Challacombe, J. F., Pillai, S., & Kuske, C. R. (2017). Shared features of cryptic plasmids from environmental and pathogenic Francisella species. PLOS ONE, 12(8), e0183554. 10.1371/journal.pone.0183554

Chambers, E. W., Hapairai, L., Peel, B. A., Bossin, H., & Dobson, S. L. (2011). Male Mating Competitiveness of a *Wolbachia*-Introgressed *Aedes polynesiensis* Strain under Semi-Field Conditions. PLOS Neglected Tropical Diseases, 5(8), e1271. 10.1371/journal.pntd.0001271

Clements, A.N. (1992). The Biology of Mosquitoes (vol. 1). CABI Publishing

Dedrick, R. M., Aull, H. G., Jacobs-Sera, D., Garlena, R. A., Russell, D. A., Smith, B. E., Mahalingam, V., Abad, L., Gauthier, C. H., & Hatfull, G. F. (2021). The Prophage and Plasmid Mobilome as a Likely Driver of Mycobacterium abscessus Diversity. mBio, 12(2), 10.1128/mbio.03441-20. https://doi.org/10.1128/mbio.03441-20

Del Solar, G., & Espinosa, M. (2000). Plasmid copy number control : An ever-growing story. Molecular Microbiology, 37(3), 492–500. 10.1046/j.1365-2958.2000.02005.x

del Solar, G., Giraldo, R., Ruiz-Echevarría, Espinosa, M., & -OrejasAUTHORS. (1998). Replication and Control of Circular Bacterial Plasmids. Microbiology and Molecular Biology Reviews, 62(2), 434–464. 10.1128/mmbr.62.2.434-464.1998

Dobson, S. L., Bourtzis, K., Braig, H. R., Jones, B. F., Zhou, W., Rousset, F., & O’Neill, S. L. (1999). *Wolbachia* infections are distributed throughout insect somatic and germ line tissues. Insect Biochemistry and Molecular Biology, 29(2), 153–160. 10.1016/S0965-1748(98)00119-2

Dumas, E., Atyame, C. M., Milesi, P., Fonseca, D. M., Shaikevich, E. V., Unal, S., Makoundou, P., Weill, M., & Duron, O. (2013). Population structure of *Wolbachia* and cytoplasmic introgression in a complex of mosquito species. BMC Evolutionary Biology, 13(1), 181. 10.1186/1471-2148-13-181

Duron, O., Fort, P., & Weill, M. (2007). Influence of aging on cytoplasmic incompatibility, sperm modification and *Wolbachia* density in *Culex pipiens* mosquitoes. Heredity, 98(6), 368–374. 10.1038/sj.hdy.6800948

Dutton, T. J., & Sinkins, S. P. (2004). Strain-specific quantification of *Wolbachia* density in *Aedes albopictus* and effects of larval rearing conditions. Insect Molecular Biology, 13(3), 317–322. 10.1111/j.0962-1075.2004.00490.x

Fallon, A. M. (2021). Growth and Maintenance of *Wolbachia* in Insect Cell Lines. Insects, 12(8), 706. 10.3390/insects12080706

Fitzgerald, S. F., Lupolova, N., Shaaban, S., Dallman, T. J., Greig, D., Allison, L., Tongue, S. C., Evans, J., Henry, M. K., McNeilly, T. N., Bono, J. L., & Gally, D. L. (2021). Genome structural variation in *Escherichia coli* O157:H7. Microbial Genomics, 7(11), 000682. 10.1099/mgen.0.000682

Fogarty, E. C., Schechter, M. S., Lolans, K., Sheahan, M. L., Veseli, I., Moore, R. M., Kiefl, E., Moody, T., Rice, P. A., Yu, M. K., Mimee, M., Chang, E. B., Ruscheweyh, H.-J., Sunagawa, S., Mclellan, S. L., Willis, A. D., Comstock, L. E., & Eren, A. M. (2024). A cryptic plasmid is among the most numerous genetic elements in the human gut. Cell, 187(5), 1206–1222.e16. 10.1016/j.cell.2024.01.039

Frost, L. S., Leplae, R., Summers, A. O., & Toussaint, A. (2005). Mobile genetic elements : The agents of open source evolution. Nature Reviews Microbiology, 3(9), 722–732. 10.1038/nrmicro1235

Gerth, M., Gansauge, M.-T., Weigert, A., & Bleidorn, C. (2014). Phylogenomic analyses uncover origin and spread of the *Wolbachia* pandemic. Nature Communications, 5(1), 5117. 10.1038/ncomms6117

Ghousein, A., Tutagata, J., Schrieke, H., Etienne, M., Chaumeau, V., Boyer, S., Pages, N., Roiz, D., Eren, A. M., Cambray, G., & Reveillaud, J. (2023). pWCP is a widely distributed and highly conserved *Wolbachia* plasmid in *Culex pipiens* and *Culex quinquefasciatus* mosquitoes worldwide. ISME Communications, 3(1), Article 1. 10.1038/s43705-023-00248-2

Hague, M. T. J., Wheeler, T. B., & Cooper, B. S. (2024). Comparative analysis of Wolbachia maternal transmission and localization in host ovaries. Communications Biology, 7(1), 1–12. 10.1038/s42003-024-06431-y

Hartig, F. (2016). *DHARMa : Residual Diagnostics for Hierarchical (Multi-Level / Mixed) Regression Models* (p. 0.4.6) [Jeu de données]. 10.32614/CRAN.package.DHARMa

Hertig, M., & Wolbach, S. B. (1924). Studies on Rickettsia-Like Micro-Organisms in Insects. The Journal of Medical Research, 44(3), 329–374.7.

Jeyaprakash, A., & Hoy, M. A. (2000). Long PCR improves *Wolbachia* DNA amplification : *Wsp* sequences found in 76% of sixty-three arthropod species. Insect Molecular Biology, 9(4), 393–405. 10.1046/j.1365-2583.2000.00203.x

Kaur, R., Shropshire, J. D., Cross, K. L., Leigh, B., Mansueto, A. J., Stewart, V., Bordenstein, S. R., & Bordenstein, S. R. (2021). Living in the endosymbiotic world of Wolbachia : A centennial review. Cell Host & Microbe, 29(6), 879–893. 10.1016/j.chom.2021.03.006

Kazlauskas, D., Varsani, A., Koonin, E. V., & Krupovic, M. (2019). Multiple origins of prokaryotic and eukaryotic single-stranded DNA viruses from bacterial and archaeal plasmids. Nature Communications, 10(1), 3425. 10.1038/s41467-019-11433-0

LePage, D. P., Metcalf, J. A., Bordenstein, S. R., On, J., Perlmutter, J. I., Shropshire, J. D., Layton, E. M., Funkhouser-Jones, L. J., Beckmann, J. F., & Bordenstein, S. R. (2017). Prophage WO genes recapitulate and enhance Wolbachia-induced cytoplasmic incompatibility. Nature, 543(7644), 243–247. 10.1038/nature21391

Lerminiaux, N. A., & Cameron, A. D. S. (2019). Horizontal transfer of antibiotic resistance genes in clinical environments. Canadian Journal of Microbiology, 65(1), 34–44. 10.1139/cjm-2018-0275

Liang, X., Liu, J., Bian, G., & Xi, Z. (2020). *Wolbachia* Inter-Strain Competition and Inhibition of Expression of Cytoplasmic Incompatibility in Mosquito. Frontiers in Microbiology, 11. 10.3389/fmicb.2020.01638

Lüdecke, D., Ben-Shachar, M. S., Patil, I., Waggoner, P., & Makowski, D. (2021). performance : An R Package for Assessment, Comparison and Testing of Statistical Models. Journal of Open Source Software, 6(60), 3139. 10.21105/joss.03139

Marimuthu, K., Venkatachalam, I., Koh, V., Harbarth, S., Perencevich, E., Cherng, B. P. Z., Fong, R. K. C., Pada, S. K., Ooi, S. T., Smitasin, N., Thoon, K. C., Tambyah, P. A., Hsu, L. Y., Koh, T. H., De, P. P., Tan, T. Y., Chan, D., Deepak, R. N., Tee, N. W. S., … Ng, O. T. (2022). Whole genome sequencing reveals hidden transmission of carbapenemase-producing Enterobacterales. Nature Communications, 13(1), 3052. 10.1038/s41467-022-30637-5

Martinez, J., Ant, T. H., Murdochy, S. M., Tong, L., Filipe, A. da S., & Sinkins, S. P. (2022). Genome sequencing and comparative analysis of *Wolbachia* strain *w*AlbA reveals *Wolbachia*-associated plasmids are common. PLOS Genetics, 18(9), e1010406. 10.1371/journal.pgen.1010406

Mavrich, T. N., & Hatfull, G. F. (2017). Bacteriophage evolution differs by host, lifestyle and genome. Nature Microbiology, 2(9), 1–9. 10.1038/nmicrobiol.2017.112

Moran, N. A., McCutcheon, J. P., & Nakabachi, A. (2008). Genomics and Evolution of Heritable Bacterial Symbionts. Annual Review of Genetics, 42(1), 165–190. 10.1146/annurev.genet.41.110306.130119

Newton, I. L. G., & Rice, D. W. (2020). The Jekyll and Hyde Symbiont : Could *Wolbachia* Be a Nutritional Mutualist? Journal of Bacteriology, 202(4). 10.1128/JB.00589-19

O’Neill, S. L. (2018). The Use of Wolbachia by the World Mosquito Program to Interrupt Transmission of Aedes aegypti Transmitted Viruses. In R. Hilgenfeld & S. G. Vasudevan (Éds.), Dengue and Zika : Control and Antiviral Treatment Strategies (p. 355–360). Springer. 10.1007/978-981-10-8727-1_24

Owashi, Y., Arai, H., Adachi-Hagimori, T., & Kageyama, D. (2024). *Rickettsia* induces strong cytoplasmic incompatibility in a predatory insect. Proceedings of the Royal Society B: Biological Sciences, 291(2027), 20240680. 10.1098/rspb.2024.0680

Partridge, S. R., Kwong, S. M., Firth, N., & Jensen, S. O. (2018). Mobile Genetic Elements Associated with Antimicrobial Resistance. Clinical Microbiology Reviews, 31(4), 10.1128/cmr.00088-17. https://doi.org/10.1128/cmr.00088-17

Reveillaud, J., Bordenstein, S. R., Cruaud, C., Shaiber, A., Esen, Ö. C., Weill, M., Makoundou, P., Lolans, K., Watson, A. R., Rakotoarivony, I., Bordenstein, S. R., & Eren, A. M. (2019). The *Wolbachia* mobilome in *Culex pipiens* includes a putative plasmid. Nature Communications, 10(1), 1051. 10.1038/s41467-019-08973-w

Richardson, E. J., Bacigalupe, R., Harrison, E. M., Weinert, L. A., Lycett, S., Vrieling, M., Robb, K., Hoskisson, P. A., Holden, M. T. G., Feil, E. J., Paterson, G. K., Tong, S. Y. C., Shittu, A., van Wamel, W., Aanensen, D. M., Parkhill, J., Peacock, S. J., Corander, J., Holmes, M., & Fitzgerald, J. R. (2018). Gene exchange drives the ecological success of a multi-host bacterial pathogen. Nature Ecology & Evolution, 2(9), 1468–1478. 10.1038/s41559-018-0617-0

Rouches, M. V., Xu, Y., Cortes, L. B. G., & Lambert, G. (2022). A plasmid system with tunable copy number. Nature Communications, 13(1), 3908. 10.1038/s41467-022-31422-0

Rousset, F., Braig, H. R., & O’Neill, S. L. (1999). A stable triple *Wolbachia* infection in Drosophila with nearly additive incompatibility effects. Heredity, 82(6), 620–627. 10.1046/j.1365-2540.1999.00501.x

Sengupta, M., & Austin, S. (2011). Prevalence and Significance of Plasmid Maintenance Functions in the Virulence Plasmids of Pathogenic Bacteria. Infection and Immunity, 79(7), 2502–2509. 10.1128/iai.00127-11

Shropshire, J. D., Leigh, B., & Bordenstein, S. R. (2020). Symbiont-mediated cytoplasmic incompatibility : What have we learned in 50 years? eLife, 9, e61989. 10.7554/eLife.61989

Siguier, P., Gourbeyre, E., & Chandler, M. (2014). Bacterial insertion sequences : Their genomic impact and diversity. FEMS Microbiology Reviews, 38(5), 865–891. 10.1111/1574-6976.12067

The, H. C., Thanh, D. P., Holt, K. E., Thomson, N. R., & Baker, S. (2016). The genomic signatures of Shigella evolution, adaptation and geographical spread. Nature Reviews Microbiology, 14(4), 235–250. 10.1038/nrmicro.2016.10

Tria, F. D. K., & Martin, W. F. (2021). Gene Duplications Are At Least 50 Times Less Frequent than Gene Transfers in Prokaryotic Genomes. Genome Biology and Evolution, 13(10), evab224. 10.1093/gbe/evab224

Unckless, R. L., Boelio, L. M., Herren, J. K., & Jaenike, J. (2009). *Wolbachia* as populations within individual insects : Causes and consequences of density variation in natural populations. Proceedings of the Royal Society B: Biological Sciences, 276(1668), 2805–2811. 10.1098/rspb.2009.0287

Viñuelas, J., Febvay, G., Duport, G., Colella, S., Fayard, J., Charles, H., Rahbé, Y., & Calevro, F. (2011). Multimodal dynamic response of the *Buchnera aphidicola* pLeu plasmid to variations in leucine demand of its host, the pea aphid *Acyrthosiphon pisum*. Molecular Microbiology, 81(5), 1271–1285. 10.1111/j.1365-2958.2011.07760.x

Weill, M., Berticat, C., Raymond, M., & Chevillon, C. (2000). Quantitative Polymerase Chain Reaction to Estimate the Number of Amplified Esterase Genes in Insecticide-Resistant Mosquitoes. Analytical Biochemistry, 285(2), 267–270. 10.1006/abio.2000.4781

Wein, T., & Dagan, T. (2020). Plasmid evolution. Current Biology, 30(19), R1158–R1163. 10.1016/j.cub.2020.07.003

Weisberg, A. J., & Chang, J. H. (2023). Mobile Genetic Element Flexibility as an Underlying Principle to Bacterial Evolution. Annual Review of Microbiology, 77(Volume 77, 2023), 603–624. 10.1146/annurev-micro-032521-022006

Werren, J. H., Baldo, L., & Clark, M. E. (2008). *Wolbachia* : Master manipulators of invertebrate biology. Nature Reviews Microbiology, 6(10), 741–751. 10.1038/nrmicro1969

Yarmolinsky, M. B. (1995). Programmed Cell Death in Bacterial Populations. Science, 267(5199), 836–837. 10.1126/science.7846528

Yu, M. K., Fogarty, E. C., & Eren, A. M. (2024). Diverse plasmid systems and their ecology across human gut metagenomes revealed by PlasX and MobMess. Nature Microbiology, 9(3), 830–847. 10.1038/s41564-024-01610-3

Zhou, W., Rousset, F., & O’Neill, S. (1998). Phylogeny and PCR–based classification of Wolbachia strains using wsp gene sequences. Proceedings of the Royal Society of London. Series B: Biological Sciences, 265(1395), 509–515. 10.1098/rspb.1998.0324

